# Classification of human white blood cells using machine learning for stain-free imaging flow cytometry

**DOI:** 10.1101/680975

**Authors:** Maxim Lippeveld, Carly Knill, Emma Ladlow, Andrew Fuller, Louise J Michaelis, Yvan Saeys, Andrew Filby, Daniel Peralta

## Abstract

Imaging flow cytometry (IFC) produces up to 12 different information-rich images of single cells at a throughput of 5000 cells per second. Yet often, cell populations are still studied using manual gating, a technique that has several drawbacks. Firstly, it is hard to reproduce. Secondly, it is subjective and biased. And thirdly, it is time-consuming for large experiments. Therefore, it would be advantageous to replace manual gating with an automated process, which could be based on stain-free measurements originating from the brightfield and darkfield image channels. To realise this potential, advanced data analysis methods are required, in particular, machine learning. Previous works have successfully tested this approach on cell cycle phase classification with both a classical machine learning approach based on manually engineered features, and a deep learning approach. In this work, we compare both approaches extensively on the complex problem of white blood cell classification. Four human whole blood samples were assayed on an ImageStream-X MK II imaging flow cytometer. Two samples were stained for the identification of 8 white blood cell types, while two other sample sets were stained for the identification of resting and active eosinophils. For both datasets, four machine learning classifiers were evaluated on stain-free imagery using stratified 5-fold cross-validation. On the white blood cell dataset the best obtained results were 0.776 and 0.697 balanced accuracy for classical machine learning and deep learning, respectively. On the eosinophil dataset this was 0.866 and 0.867 balanced accuracy. From the experiments we conclude that classifying distinct cell types based on only stain-free images is possible with these techniques. However, both approaches did not always succeed in making reliable cell subtype classifications. Also, depending on the cell type, we find that even though the deep learning approach requires less expert input, it performs on par with a classical approach.

## Introduction

Imaging flow cytometry (IFC) produces up to 12 spectrally distinct, information-rich images of single cells at a throughput of up to 5000 cells per second with a resolution of 0.25 μm per pixel (60x magnification) (1). This includes at least two label-free image channels produced by transmitted (bright-field) and scattered light (dark-field). These characteristics make IFC an ideal candidate for in-depth analyses of cell populations as an approach to unlock the inherent heterogeneity contained within all biological systems. For example, IFC has been used to detect rare circulating endothelial cells, which have been correlated with various disease states when present in elevated levels (2). Furthermore, it has also been applied to replace the use of manual microscopy for the in vitro micronucleus assay used to study geno- and cytotoxicity. The use of IFC allowed for more automation and informative visualizations (3). IFC has also been used to study the partitioning of molecules across the plane of cell division in a statistically robust manner (4, 5, 6).

In this and most other IFC research, cell populations are studied with manual gating on numerical features extracted from the IFC data by specialized software. With manual gating, cells are hierarchically divided into sub-populations by setting boundaries, or *gates*, on 2D scatter-plots of cell measurements. These measurements are usually a combination of fluorescence intensities derived from a targeted probe against a feature of a cell of interest, and morphological characteristics derived from the cell images. Although this approach has led to numerous insights into cell population heterogeneity (7), it has some serious drawbacks, mainly:

i. manual gating is hard to reproduce,
ii. manual gating is subjective and biased,
iii. and manual gating is time-consuming for large experiments (8).

Analysing IFC data with manual gating limits the potential of the information-rich, spatially registered data it provides. This is true because gating is done on 2D scatter plots, which allow only two features to be viewed at once, whereas an approach combining a multitude of features can reveal much more intricate patterns in the same data.

Manual gating is an expert-driven process, which introduces two main sources of operator bias. First, gates set on the scatter plots are highly subjective, and can therefore differ significantly between operators. The second source is specific to IFC. The choice of which features to compute, and on which area of interest in the image (referred to as *mask*) to compute them, greatly influence downstream analysis of the data. The operator skill is again an important factor of variability (9).

Fluorescent stains have drawbacks as well. Firstly, there are potential detrimental effects on the cells under study influencing achieved results (10, 11). Secondly, usually several stains are required to precisely identify a cell (12), making the experimental workflow labor-intensive and slower. Because of these reasons *stain-free* experiments have become of particular interest in the bio-imaging field over the last decade (13, 14, 15).

A potential solution for overcoming these drawbacks is to automate the gating process with machine learning (ML), and do this with only stain-free measurements. This approach (1) combines all available features by using complex ML models, (2) limits operator bias through automation, and (3) potentially obsoletes fluorescent staining by using only stain-free measurements to perform cell classification.

Previous work has explored this approach, and formulated it as a machine learning image classification task, where the goal is to train a supervised classification model on features computed on stain-free imagery. Hennig et al. (16) developed an open-source solution, which uses the software package CellProfiler (17) to extract image features from stain-free cell imagery, and classical supervised machine learning to classify the cells in sub-populations. They were able to classify Jurkat cells into 5 phases of the cell cycle.

Another example is the work by Eulenberg et al. (18), who developed a deep learning (DL) model, termed *DeepFlow*. It is able to reconstruct the cell cycle of Jurkat cells, as well as to study the disease progression of diabetic retinopathy. DeepFlow is a convolutional neural network (CNN), which autonomously extracts relevant features from input images to perform a classification, eliminating the requirement for specialized tools to extract features. Deep learning is currently widely used in image classification, and is increasingly being adopted in image cytometry.

In this work we contribute to the previous work by extensively comparing both classical ML and DL, testing out two models per approach. We test classification performance of all models on two high quality white blood cell datasets from healthy human whole blood samples, acquired on an ImageStream^X^ MK-II platform. Unlike the work mentioned above, these datasets do not focus on the cell cycle of Jurkat cells, but on the identification of various types of white blood cells. In addition, the first dataset contains specific cell subtypes (for example, CD4+ and CD8+ T-cells). In the second dataset, active and resting eosinophils are identified. Activation state is of importance as elevated eosinophil activation is linked with allergic disease and intrinsic asthma for example (19, 20). The stain-free classification of these more subtle cell types has not been attempted in previous work, and is challenging as differences in their stain-free measurements are expected to be less pronounced. Furthermore, we also compare the feature spaces used for classification with dimensionality reduction techniques.

## Materials and Methods

### Datasets

All datasets are acquired from human blood samples. Ethical approval to obtain blood from healthy volunteers was granted by the County Durham and Tees Valley Research Ethics Committee (12/NE/0121).

#### White blood cells (WBC)

This dataset contains the measurement results from white blood cells from 2 whole blood samples collected into citrate buffer. For phenotyping experiments, 500 μl of whole blood was placed in a 15 ml falcon tube, so that approximately 2 × 10^6^ WBCs were stained with the following antibody cocktail: CD15 FITC (BD, cat no: 332778, clone MMA, 5 μl per test), Siglec8 Pe (Biolegend, cat no: 347104, clone 7C9), CD14 PeCF594 (BD, cat no: 562334, clone M*ϕ*9, 5 μl per test), CD19 PerCP-CY5.5 (BD, cat no 340951, clone SJ25C1, 20 μl per test), CD3 BV421 (BD, cat no: 562426, clone UCHT1, 5μl per test), CD45 V500 (BD, cat no: 647450, clone 2D1, 5 μl per test), CD4 BV605 (BD, cat no: 562658, clone RPA-T4, 5 μl per test), CD56 APC (BD, cat no: 341025, clone NCAM16.2, 5μl per test) and CD8 APC-CY7 (BD, cat no: 557834, clone SK1, 5μl test). Whole blood was incubated with the staining cocktail for 1 hour on ice after which red blood cell (RBC) lysis was performed by the addition of 4.5 ml of 1x BD FACS lysis solution (cat no: 349202) prepared from a 10x stock in reagent grade water (SIGMA, cat no: W4502). Lysis was carried out for 10 minutes at RT in the dark. Samples were then spun down at 500 g for 5 minutes and washed twice in 50 ml of wash buffer (PBS + 2% FBS). Samples were re-suspended in a final volume of 60 μl of wash buffer and transferred to 1.5 ml Eppendorf tubes for acquisition.

This panel allowed for the identification of eight different white blood cell types: CD4+/CD8+ T-cells, neutrophils, monocytes, B-cells, CD56+ NKT-cells, other NKT-cells and eosinophils. The data was analysed by expert annotation with hierarchical manual gating using the fluorescence marker information, akin to phenotyping by conventional flow cytometry. See Figure 3 for an overview of the gating process.

The dataset is imbalanced: it contains 17358 CD4+ T-cells, 8 022 CD8+ T-cells, 59 034 CD15+ neutrophils, 2 655 monocytes, 4 256 CD19+ B-cells, 2 214 CD56+ NKT-cells, 1318 other NKT-cells, and 3156 eosinophils.

#### Eosinophils (EOS)

This dataset contains the measurement results from white blood cells from 2 whole blood samples collected into Heparin buffer. For eosinophil activation experiments, 1 ml of whole blood was transferred to 15 ml Falcon tubes, one for each of the following conditions: 1) Ex-vivo control that was kept on ice for the duration of stimulation, 2) 20 minute stimulation, 3) 40 minute stimulation, 4) 60 minute stimulation, 5) unstimulated control, incubated for the duration of stimulation. In the first instance, stimulations were performed using PMA/Ionomycin (eBiosciences Cell Stimulation Cocktail, cat no: 00-4970-03) at a 1x working concentration. In order to ensure all incubations ended at the same time, the 60 minute stimulation was started first, then the 40 minute stimulation, 20 minutes later, and finally the 20 minute stimulation a further 20 minutes later. At the end of the stimulation period, the samples were divided in two 15ml Falcon tubes (500μl in each). One sample set was stained with the following antibody cocktail: Siglec8 Pe (Biolegend, 5 μl per test), CCR3/CD192 BV421 (Biolegend, cat no: 310714, clone 5E8, 5μl per test), CD69 APC (BD, cat no: 555533, clone FN50, 20μl per test), and CD11b (Biolegend, cat no: 101226, clone M1/70, 5μl per test).

The other set of the samples were left unstained to control for the effects of antibody labelling. Samples were incubated for 1 hour on ice after which RBC lysis was performed as described for the WBC panel. Again samples were washed 2 times in wash buffer and finally resupsended in 60 μl of the same for transfer into 1.5 ml Eppendorf tubes prior to acquisition.

This panel allowed for the identification of eosinophils in their active and resting state. The data was analysed by expert annotation with hierarchical manual gating using the fluorescence marker information. See Figure 3 for an overview of the gating process.

The dataset is imbalanced: it contains 1291 active and 2 595 resting eosinophils, and 186 671 non-eosinophils.

### ImageStream^x^ MKII imaging flow cytometer

IFC was performed using a fully ASSISTED-calibrated ImageStream^x^ MKII (Luminex Corporation, Seattle, WA, USA) system with the following configuration: 100 mW 488 nm blue laser, 120 mW 450 nm violet laser and a 642 nm 150 mW red laser. In all cases, maximum excitation laser powers were employed in order to achieve best signal to noise without any pixel saturation (raw max pixel values below 4096). Bright-field imagery was collected using an LED array with wavelengths of 420 nm to 480 nm in channel 1 and 570 nm to 595 nm in channel 9. Side scatter was collected in channel 6 using a dedicated 758 nm laser, again set to maximize signal and avoid saturation. FITC emission was collected in CH2, Pe in CH3, PeCF594 in CH4, PerCP-CY5.5 in CH5, BV421 in CH7, V500 in CH8, BV605 in CH10, APC in CH11 and APC-CY7 in CH12. The images were acquired in the highest sensitivity mode and using the 60x magnification. For the stained and unstained samples, each were acquired with excitation lasers on and off to control for any potential for residual fluorescence spill over in to label-free channels even after spectral correction. Spectral correction was performed using the built-in wizard in the IDEAS analysis software package. Antibody capture beads (ABC total capture beads, Thermo scientific, cat no: A10513) were used to prepare single stained controls by adding 1 drop of positive, 1 drop of negative and then 1 test amount of each individual antibody per tube. These controls were acquired with the bright-field and side scatter illumination turned off in order to generate the spillover matrix. This matrix was then applied to fully stained samples.

### Classification models

Models in this work were divided into two categories: i) models which took pre-computed manually engineered features as input, and ii) models which took images as input. The latter type of models automatically learned and extracted required features from the input images. These models, usually convolutional neural networks, have been applied successfully in many image classification tasks (21, 22, 23). Using models with automatically engineered features further reduced the influence of expert knowledge on the gating process, at the cost of an increase in the computational effort required to train the model.

In total four classification models were tested, two per model category. For the first model category, referred to as classical ML, we tested a random forest (RF) (24) and gradient boosting (GB) classifier (25). Both models are ensembles of weak decision trees, which are widely used and successful in classification settings (26, 27). The number of used trees was set to 500 for random forest, and 100 for gradient boosting. All other hyper-parameters were kept at default values provided by the scikit-learn library (28).

For the second model type we tested two DL convolutional neural network (CNN) architectures: ResNet18 (RN) (22) and DeepFlow (DF) (29).

ResNet is a state-of-the-art architecture in image classification. It eases optimization of the network’s weights by reformulating convolutional layers as learning residual functions, with reference to the layer inputs. ResNet has obtained the first place on the ImageNet Large-Scale Visual Recognition Challenge 2015, an important competition in the field of computer vision (22). In this work a variant of 18 layers deep is used.

DeepFlow is an adaptation of the Inception architecture (30), optimized for classification of imaging flow cytometry data. It was previously applied to reconstruct the cell cycle of Jurkat cells, as well as to study the disease progression of diabetic retinopathy, using stain-free IFC imagery.

Both DL models were implemented in the Keras-Tensorflow Python library (version 1.13.1) (31). DeepFlow implementation was based on code from the DeepFlow Github Repository^1^. ResNet18 implementation was taken from the Keras-ResNet Github Repository^2^. The models were trained for 100 epochs with the Adam optimization algorithm (32). Learning rate was set to 10^−5^. L2 regularization was applied with a weight of 10^−4^ for ResNet18, and 5 × 10^−4^ for DeepFlow. All code used to generate results in this work is publicly available on Github^3^.

### Data preparation

To run the experiments we needed all stain-free imagery and accompanying masks, features computed on stain-free imagery, and ground truth cell type labels.

After spectral compensation (see Supplementary Table 3 for compensation matrix), the IDEAS software produces one compensated image file (CIF) per biological sample, containing compensated imagery for all channels, as well as the accompanying masks. These masks indicate the area of the image containing only the pixels of interest in a certain channel. In the case of brightfield images, the mask typically encloses the entire cell. By masking the images we avoid influence of background noise or irrelevant information surrounding the cell (33) (see Figure 1).

**Figure 1:**
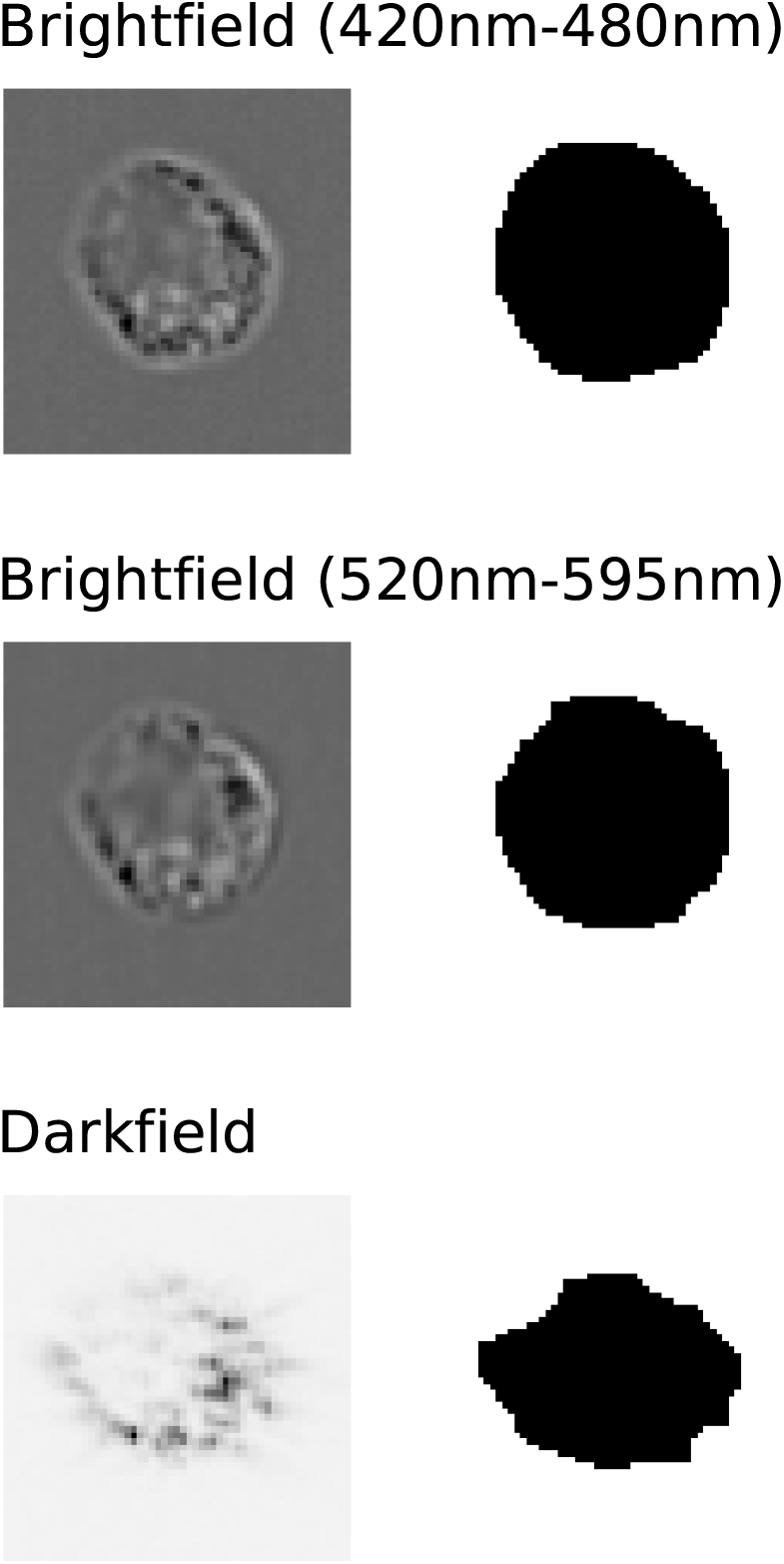
The 3 stain-free images used in this work acquired with the Amnis ImageStream-X MKII instrument capture morphological information about the cell. Images for each channel and accompanying masks for one random cell are shown, respectively in the left and right column. Masks are computed by IDEAS.

**Figure 2:**
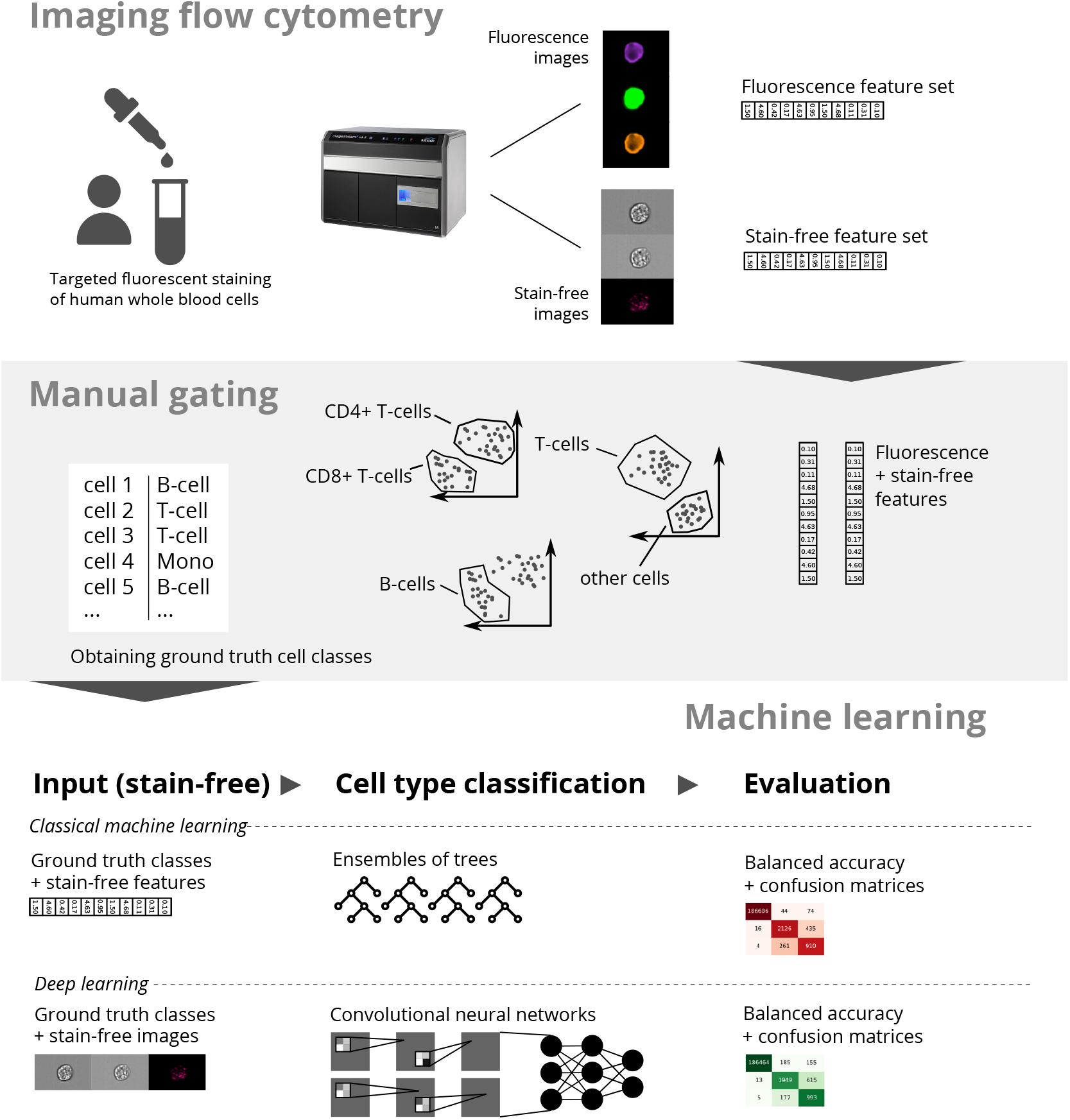
Machine learning enables cell classification based on stain-free imaging flow cytometry imagery. White blood cells from healthy humans are imaged by an imaging flow cytometer, in our case the ImageStream^x^ MK-II. Features are extracted from stain-free and fluorescence imagery. These features are used in a manual gating procedure to obtain ground truth data. This ground truth and accompanying stain-free images and features are used to train classical machine learning and deep learning models to perform cell classification.

In order to work efficiently with these images and masks, they were read and decoded from the CIF format using the Python Bio-Formats library (34), and saved to an HDF5 dataset. To perform this step a custom command line tool was written in Python. By decoding the images once and saving them in decoded form we can significantly speed up the training of the neural network, as decoding images is a costly operation. The code for this tool is made publicly available on GitHub^4^.

The first feature set was computed in IDEAS. The software computes 76 base features per image channel/mask, divided into five categories: size, location, shape, texture, signal strength and singal strength. Features are computed on the masked channel images. More details can be found in the IDEAS documentation (33).

### Data augmentation

Many classifiers, including the ones used in this work, are sensitive to class imbalance (35). Therefore, we augmented the datasets before training in order to balance the class occurrence frequencies. For the two classical ML algorithms considered in this paper, this was done by randomly oversampling minority classes.

An image-based data augmentation approach is required for the DeepFlow classifier. As is commonly done for convolutional neural networks, we apply random *label-saving* image transformations to existing training instances, supplementing the data manifold (21, 36). We used random horizontal or vertical flips, rotations, and translations.

### Model validation

To validate the classification performance of the trained models, a stratified 5-fold cross validation (CV) strategy is used. For each fold, training data was augmented to balance the class occurrence frequencies, and used to train a model. The model was then validated using non-augmented instances from the validation set. The predictions made on the validation data were summarized in a confusion matrix per fold. The obtained matrices were summed together, giving one representative confusion matrix per CV experiment.

Together with the confusion matrix, the *balanced accuracy* was reported. The balanced accuracy is the arithmetic mean of class-specific accuracies. It can be computed from the confusion matrix, and is formalized as follows:

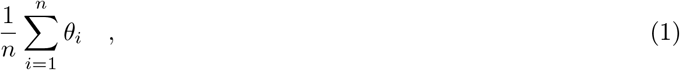

where *θ_i_* is the class-specific accuracy, and *n* is the number of classes. The balanced accuracy is suited for imbalanced validation sets, as it does not suffer from the accuracy paradox. This means it won’t favor a classifier that exploits class imbalance by biasing towards the majority class (37).

### Visualizing feature spaces

Dimensionality reduction techniques can be used to give an insight into high-dimensional spaces, by projecting them onto a low-dimensional space. In this work we applied Uniform Manifold Approximation and Projection (UMAP) (38) on the high-dimensional, manually engineered feature space exported from IDEAS, and the feature space automatically learned by the DeepFlow CNN.

UMAP provides scalable dimensionality reduction, which preserves global and local structures of the highdimensional input space. It does so by converting high- and low-dimensional representations of the input to topological representations, and then minimizing the cross-entropy between them. We choose this method over others such as t-SNE, due to its scalability and wide-spread application in bioinformatics (38).

The feature space learned by a CNN is encoded by the intermediate activation pattern following the last convolutional layer of the network, referred to as the *code*. The code is the representation of an input image, which is fed to the fully-connected layers of the CNN that perform the actual classification. The codes are extracted from the network by forward-propagating images through it, and recording their corresponding codes.

All ML experiments were run on a 12-core machine, with an Intel Xeon CPU (model E5-1650 v2) running at 3.50GHz. The machine has 64 GB of RAM. DL experiments were run on an NVIDIA Titan X GPU with 12 GB of VRAM. Code used for extracting imagery and masks from the CIFs, and for training and validation of the models is made public on GitHub at https://github.com/saeyslab.

## Results

We started by setting a baseline classification performance on the EOS and WBC datasets, using well established models, trained on expert-driven, manually engineered features. We then applied DL to the same classification tasks and found that they achieved baseline performance for the EOS dataset, but not for the WBC dataset.

### Classifying cell types with manually engineered features

Both classical ML classifiers were able to classify cell types based on manually engineered features (see Figures 5 and 6). For the WBC dataset, especially neutrophils, monocytes and eosinophils were accurately classified by both classifiers with recalls respectively higher than 0.974, 0.957 and 0.965 for all classifiers. For the EOS dataset, separation between non-eosinophils and eosinophils was very accurate, with recalls respectively higher then 0.998 and 0.990 (computed by treating active and resting eosinophils as one class).

**Figure 3:**
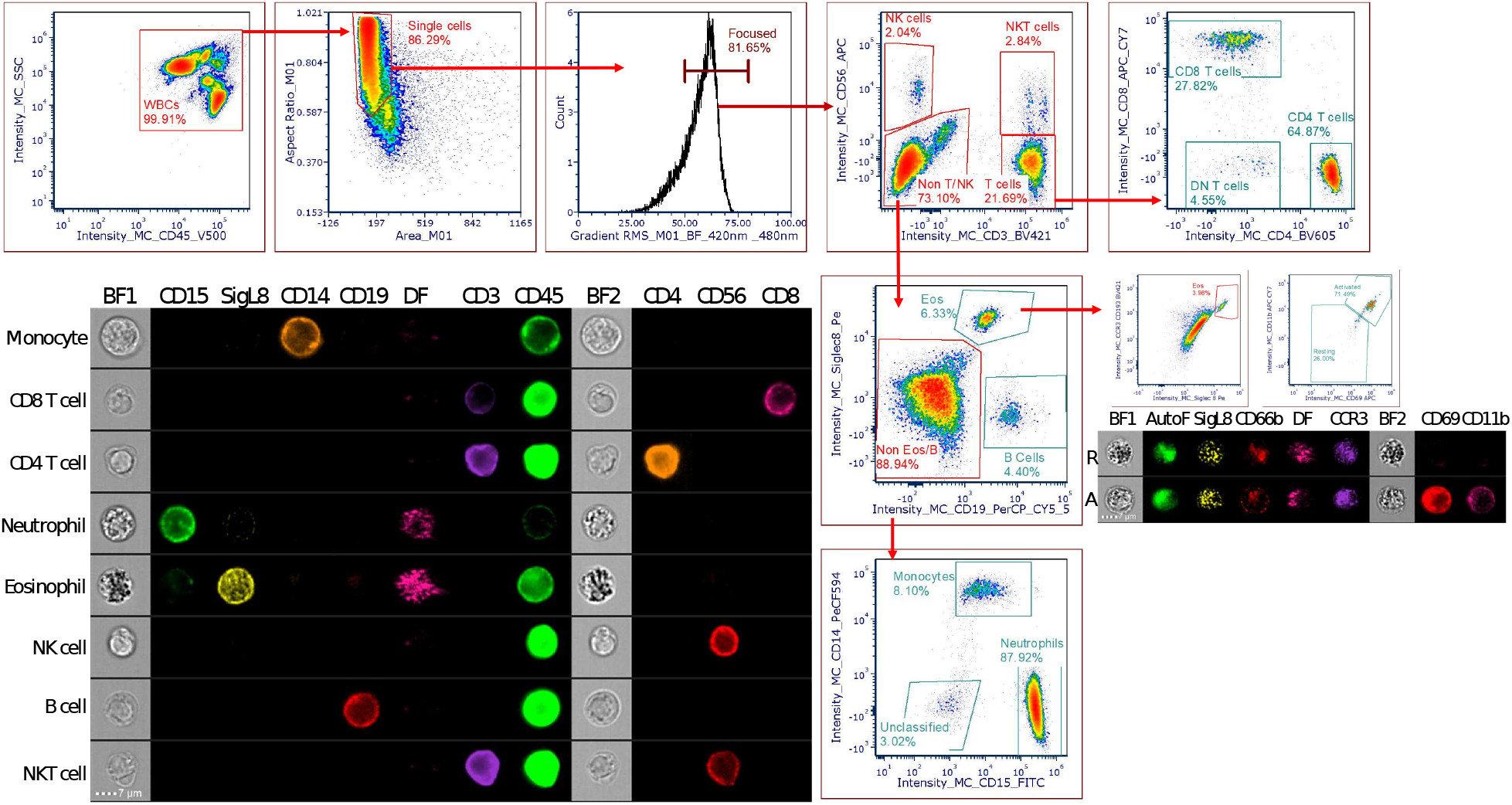
The “ground truth” gating strategy based on fluorescence antibody information. Briefly, White Blood Cells (WBCs) were gated based on CD45 V500 fluorescence and darkfield (DF). Single cells were identified based on the Area and Aspect ratio of the bright-field image in channel (BF1). Focused cells were identified based on the Gradient RMS feature (>50AU). NK, NKT and T cells were then identified based on CD3 and CD56 fluorescence with the T cell population further subdivided in to CD4 and CD8 subsets. The remaining cells were identified as either B cells (CD19 positivity), Eosinophils (Siglec 8 positivity), Monocytes (CD14 positivity) or Neutrophils (CD15 positivity). Example multi-spectral, compensated images are shown for each class of immune cell identified at 60x magnification. It should also be noted that for ML/DL, cells were always compensated and samples were pre-processed to the stage of CD45 positive, single in focus cells, as shown above.

**Figure 4:**
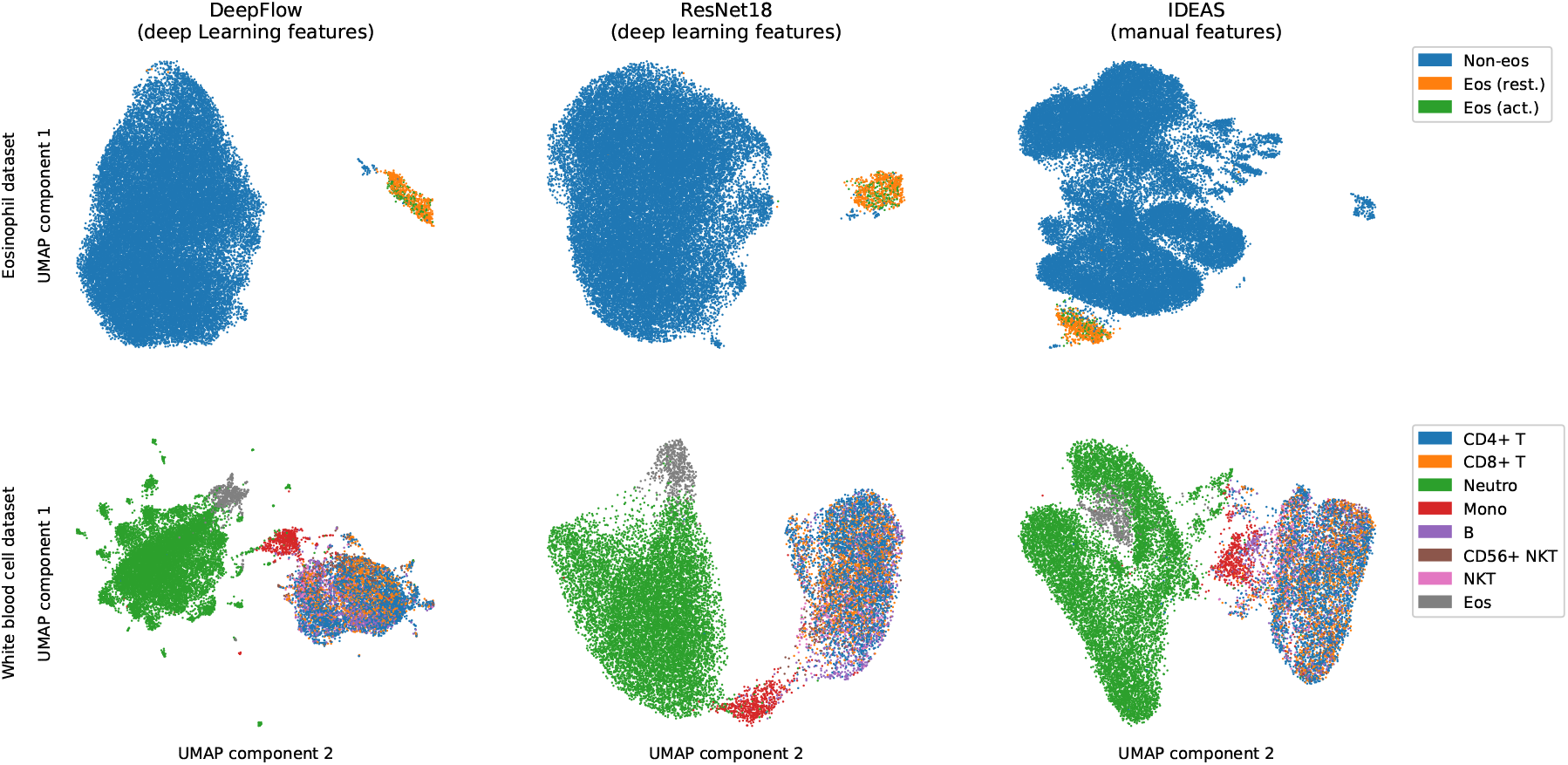
Dimensionality reduction of manually and automatically engineered feature spaces confirmed confusion in cell type classification. High-dimensional feature spaces were projected to a 2D space using Uniform Manifold Approximation and Projection. Data points were plotted in the 2D space and colored according to cell type. This revealed that cells of the same cell type cluster together. Clusters of cells types that overlapped, were also found challenging to distinguish in the classification experiments. For example, CD4+ and CD8+ T-cells overlapped significantly in the white blood cell dataset and also show high confusion in the classification. The same is seen for the active and resting eosinophils in the eosinophil dataset.

**Figure 5:**
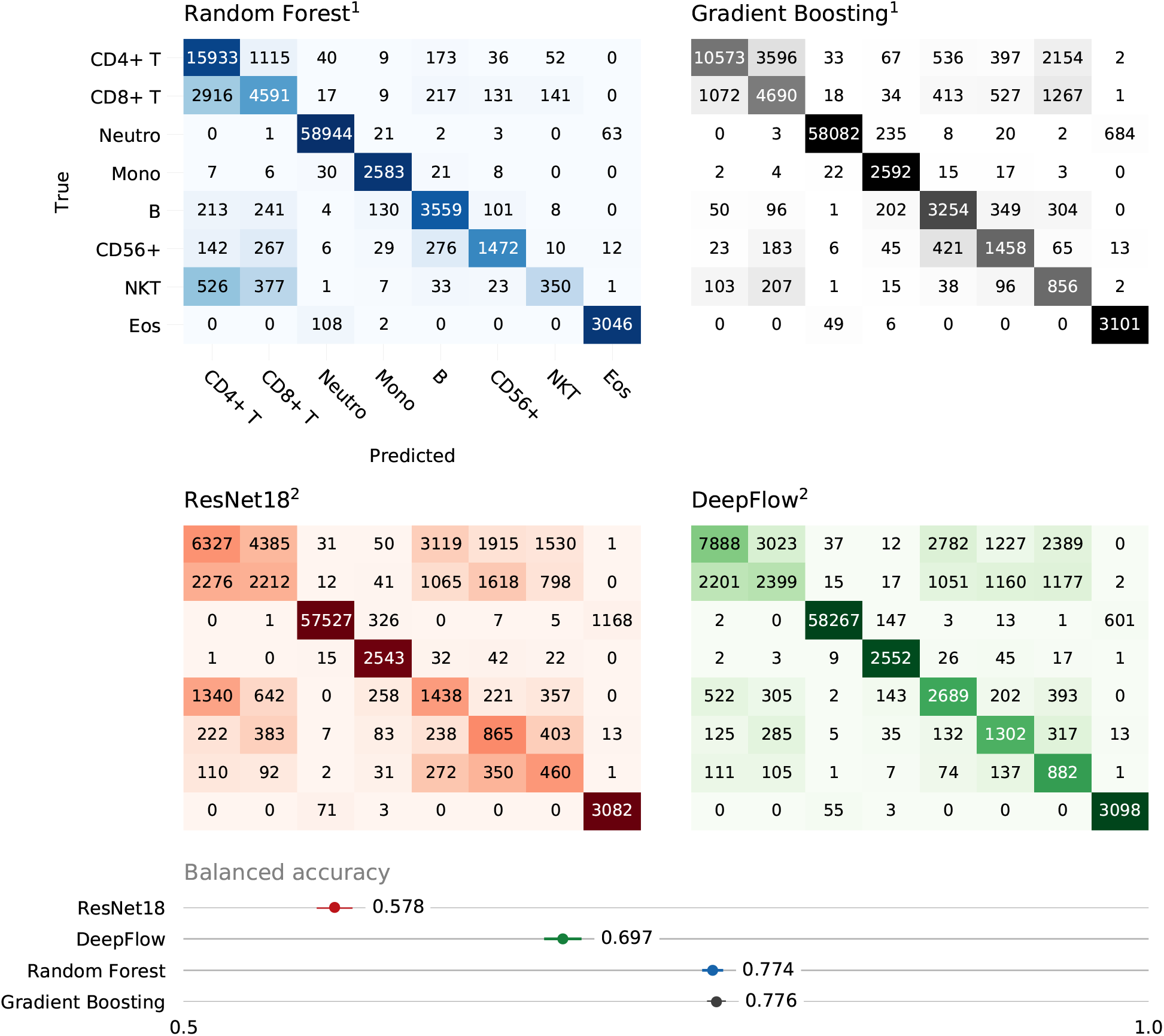
Classifiers based on manually engineered features (1) outperformed deep learning classifiers using automatically learned features (2) on the white blood cell dataset. As seen on on the confusion matrices, neutrophils, monocytes and eosinophils are consistently classified correctly by all classifiers, as seen in the confusion matrices. Confusion is present in all classifiers when subtyping T-cells or NKT-cells. Both classical machine learning classifiers behave similarly and outperform the deep learning classifiers.

**Figure 6:**
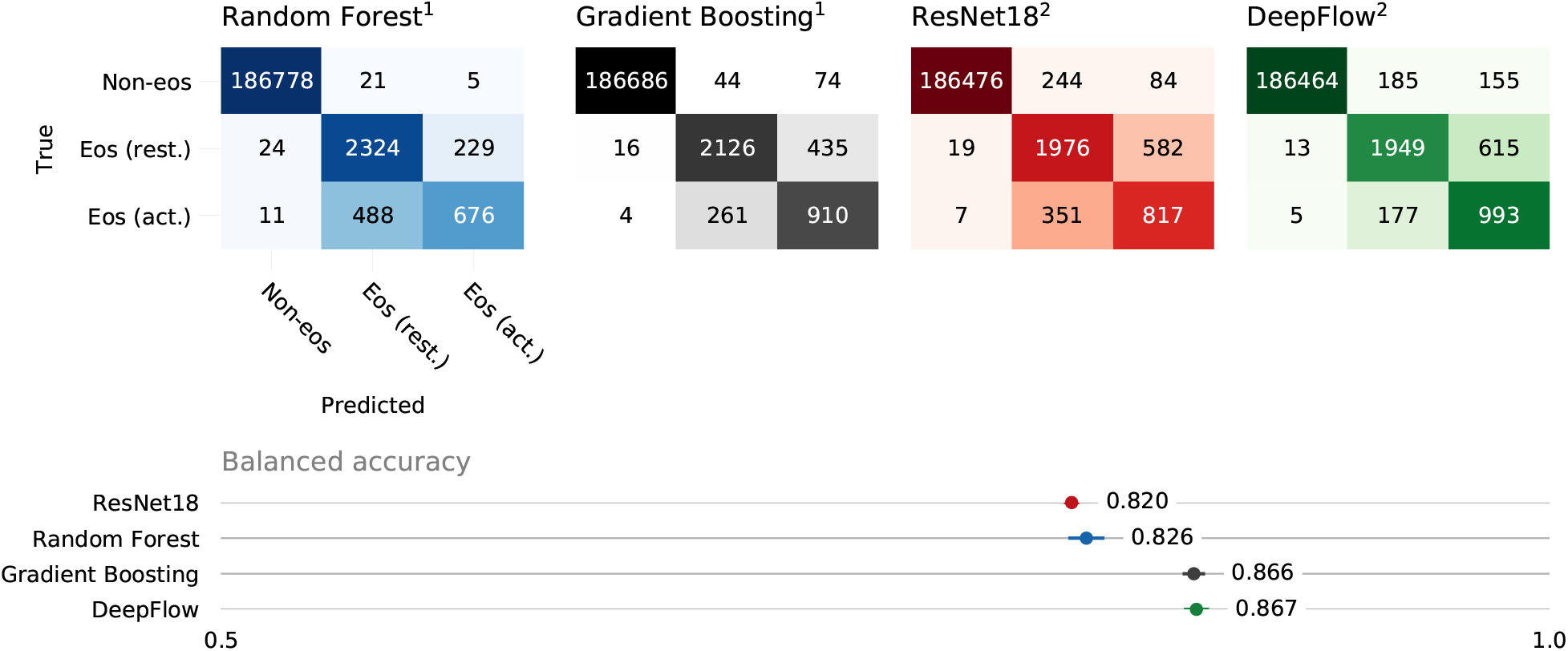
Classifiers based on automatically learned features (2) performed on par with classifiers using manually engineered features (1) on the eosinophil dataset. As seen on the confusion matrices, separation between eosinophils and non-eosinophils is consistently done correctly by all classifiers. Confusion is present in all classifiers when separating between active and resting eosinophils, especially the random forest struggled to make a good distinction. Between deep learning approaches, the DeepFlow classifier outperformed ResNet18 by a small margin.

Both classifiers struggled to reliably subtype cell types. Recalls for the subtypes are consistently lower than for other classes (see Supplementary Table 1 and 2. In the WBC dataset confusion was present between CD4+ and CD8+ T-cells for example, as well as between T-cells and NKT-cells (see Figure 5). Also in the EOS dataset confusion was present between active and resting eosinophils (see Figure 6).

We found that the GB and RF classifier behaved similarly, with a slight advantage for the GB classifier on the EOS dataset. Their respective balanced accuracies were 0.774 versus 0.776 for the WBC dataset, and 0.826 versus 0.866 for the EOS dataset (see Figure 5 and 6). The main advantage of the GB over the RF classifier was the better subtyping performance, which is clearly seen in the classification of active versus resting eosinophils, and CD4+ versus CD8+ T-cells.

### Automating feature extraction with deep learning

DL classifiers were able to autonomously extract relevant information from stain-free imagery to classify WBCs and eosinophils (see Figure 5 and 6). Their performance differed between both datasets: in the EOS dataset DeepFlow improved slightly upon the baseline performance set by both classical methods (see Figure 6). However, in the WBC dataset none of the DL classifiers reached baseline performance (DF: 0.697, versus GB: 0.775 balanced accuracy) (see Figure 5).

As with the classical models, the DL models did not reach satisfactory performance for cell subtyping. In the WBC dataset, recalls for CD4+ and CD8+ T-cells did not exceed 0.454 and 0.299, respectively. NKT-cell subtyping suffered less of a drop in recall compared to classical methods, with recall values reaching 0.588 and 0.669 for CD56+ and other NKT-cells, respectively.

Overall, the DF architecture outperformed the RN architecture. The difference was most pronounced in the WBC dataset (RN: 0.578, versus DF: 0.697 balanced accuracy). Recalls for all cell types were higher for DF. The biggest improvement over RN occurred in B-cell classification (RN: 0.391, versus DF: 0.632 recall). In the EOS dataset, the improvement from DF over RN was smaller (RN: 0.820, versus DF: 0.867 balanced accuracy).

### Comparing feature spaces with Uniform Manifold Approximation and Projection

Visualizing the feature spaces on which the classifiers are trained, provided a visual validation of classification confusion occurring between certain cell types (see Figure 4). UMAP clustered cells of similar cell types together in the low-dimensional representation. We found that clusters of cell types overlapped for the types with which classifiers struggled. For example, in the WBC dataset, confusion occurred between CD4+ and CD8+ T-cells (see Figure 5). This is clearly reflected in the UMAP visualization by the overlapping clusters of CD4+ and CD8+ T-cells, both for automatically and manually engineered features. On the other hand, accurately classified cell types, such has the eosinophils, were also well separated in the low-dimensional space. The same occured in the EOS dataset for the confusion between active and resting eosinophils (see Figure 4).

## Discussion

Manual gating in its current form has three main drawbacks: manual gating is hard to reproduce, it is subjective and biased, and it is time-consuming for large experiments. To overcome these drawbacks we extensively studied and compared several ML approaches, which only make use of stain-free information to perform automated cell classification.

In the experimental setting of this work, stains do not seem vital for classifying cell types such as monocytes or neutrophils. However, to reliably subtype cells, stains are still required. This is shown in Figure 5, and Supplementary Table 1. The best performing model, a gradient boosting classifier, has a recall higher than 0.976 for neutrophils, monocytes and eosinophils. The overall T-cell recall is 0.785, but for the CD4+ and CD8+ T-cell subtypes, recall rates drop to 0.609 and 0.585, respectively. This means that visually, based on the stain-free images, no distinction can be made between CD4+ and CD8+ T-cells. The same pattern can be observed in Figure 6 and Supplementary Table 2 for active and resting eosinophils. Other works have concluded also that classifying distinct cell types using stain-free information is possible (39), as well as classifying cell cycle phases (40). To the best of our knowledge, this contribution is the first attempt to classifying cell subtypes using purely stain-free information.

The unreliable cell subtyping might be attributed to two data-related issues: class imbalance and low image resolution. Firstly, all datasets in this work suffer from class imbalance. For example, the EOS dataset contains about 187 000 non-eosinophils, and only about 3 900 eosinophils, with a 30 vs 70% ratio of active and resting eosinophils. Especially in DL settings, large and balanced datasets are important, as overfitting occurs regularly (41). In this work we employ basic data augmentation techniques to counter this problem, which seem to have improved performance to a certain degree. Since acquiring more data is not always an option, developing or employing more advanced data augmentation techniques could improve performance.

Secondly, because of the relatively low image resolution of current imaging flow cytometers, necessary data for cell subtyping is potentially not captured in the stain-free imagery. A solution, which does not eliminate, but reduces the necessary staining, might be to design hybrid experiments. These would rely on stain-free information to identify cell types and use a limited amount of stains to further subtype cells. For example, in the WBC dataset lymphocytes (NKT-, T-, and B-cells), granulocytes (neutrophils and eosinophils), and monocytes can be accurately separated using only stain-free information with the models trained in this work. The fluorescence channels that are normally reserved for identifying these larger populations, can then be employed for reliably subtyping cells using targeted staining, therefore increasing the possible number of cell populations that can be identified with the same instrumentation.

DL approaches are able to autonomously extract relevant information from stain-free imagery, a conclusion that is supported by previous work (29). For the WBC dataset the best results are achieved by classical approaches with a fairly large margin on the DL approaches (GB: 0.775 vs DF 0.697 balanced accuracy). For the EOS dataset, however, the best result is achieved by a DL approach. We could state that in this regard DL can make cell classification based on IFC imagery a less manual, and expert-driven process. However, we must note that training and optimizing neural networks for a certain classification task is not straight-forward. It requires significant expert knowledge to overcome obstacles such as overfitting, hyper-parameter tuning, handling big data, and dealing with a shortage of, or imbalance in labelled data (18, 42). Different methods to deal with these issues, such as transfer learning or data augmentation, need to be made accessible and easy to use. Therefore, we have made our code and trained models publicly available on Github^5^. In order to increase accessibility of this solution, we consider the development of an extension for the CellProfiler software, which has gained considerable popularity in the bio-imaging field.

Another difference between automatically and manually engineered feature spaces is shown by the UMAP dimensionality reduction. It shows that the manually engineered feature set is better suited for exploring the heterogeneity of cells within a dataset. This is because the automatically engineered feature spaces from the DeepFlow and ResNet18 CNNs are only optimized to distinguish between the ground truth cell types in the dataset. On the other hand, the manually engineered feature set from IDEAS is more general. This is demonstrated by the UMAP reductions for the EOS dataset in Figure 4: the non-eosinophils are clustered in one homogeneous cluster in the DL feature space, whereas several clusters can be distinguished within the non-eosinophils in the IDEAS feature space. The IDEAS features therefore seem to capture more of the heterogeneity within this population.

File formats produced by the Amnis ImageStream platform are closed-source, and therefore unsuitable for data science applications. In previous work an approach was proposed, which requires the user to create many image montages from the images in the original CIF, using a custom script (16). These montages can then be processed by image analysis software, such as CellProfiler. This is a cumbersome and non user-friendly process. In this work we have accommodated for this inconvenience by writing a script that decodes images and masks from the original CIF, and stores them in one HDF5 dataset to be used during further processing. This way we have significantly reduced processing time of the CIFs, opening the possibility to train and test ML models on hundreds of samples. The script is publicly available on Github^6^.

In conclusion we have found that imaging flow cytometry lends itself very well to ML applications due to its information-rich data and high-throughput nature. We have shown that besides cell cycle phase classification, white blood cell type classification is also feasible, creating the potential to apply this approach in immunodeficiency diagnosis, for example. For the datasets and classification methods studied here we conclude that the limit of this classification approach currently lies at the level of the cell type.

## Supplementary Material

**Supplementary Table 1:** Classification recall obtained on the white blood cell dataset. All classifiers succeeded in classifying neutrophils, monocytes and eosinophils, but have lower performance when classifying CD4+ and CD8+ T-cells, and CD56+ and other NKT-cells.

|  |  | Random Forest |  | Gradient Boosting |  | ResNet18 |  | DeepFlow |  |
| --- | --- | --- | --- | --- | --- | --- | --- | --- | --- |
| T-cells | CD4+ | 0.96749 | 0.91791 | 0.78530 | 0.60911 | 0.59890 | 0.36450 | 0.61115 | 0.45443 |
|  | CD8+ |  | 0.57230 |  | 0.58464 |  | 0.27574 |  | 0.29905 |
| Neutrophils |  |  | 0.99848 |  | 0.98387 |  | 0.97447 |  | 0.98701 |
| Monocytes |  |  | 0.97288 |  | 0.97627 |  | 0.95782 |  | 0.96121 |
| B-cells |  |  | 0.83623 |  | 0.76457 |  | 0.33788 |  | 0.63181 |
| NKT-cells | CD56+ | 0.52521 | 0.66488 | 0.70073 | 0.65855 | 0.74690 | 0.39071 | 0.58838 | 0.58809 |
|  | Other |  | 0.26557 |  | 0.64947 |  | 0.34909 |  | 0.66924 |
| Eosinophils |  |  | 0.96515 |  | 0.98257 |  | 0.97655 |  | 0.98162 |

**Supplementary Table 2:** Classification recall obtained on the eosinophil dataset. All classifiers successfully separated eosinophils and non-eosinophils, but have lower performance when classifying resting and active eosinophils. Recalls for active and resting eosinophil classification are at an acceptable level for deep learning methods, but methods based on manually engineered features struggle with this classification.

|  |  | Random Forest |  | Gradient Boosting |  | ResNet18 |  | DeepFlow |  |
| --- | --- | --- | --- | --- | --- | --- | --- | --- | --- |
| Non-eosinophil |  |  | 0.99986 |  | 0.99937 |  | 0.99824 |  | 0.99818 |
| Eosinophil | Active | 0.99068 | 0.90178 | 0.99469 | 0.82498 | 0.99306 | 0.76674 | 0.99521 | 0.75692 |
|  | Resting |  | 0.57537 |  | 0.77397 |  | 0.69538 |  | 0.84612 |

**Supplementary Table 3:** The compensation matrix calculated by acquiring single stained beads that was applied to all multi-stained cellular samples. The columns represent the fluorochromes and the rows are the imaging channels.

|  | **BF1** | **FITC** | **Pe** | **Pe-CF594** | **PCP-CY5.5** | **DF** | **BV421** | **V500** | **BF2** | **BV605** | **APC** | **APC-CY7** |
| --- | --- | --- | --- | --- | --- | --- | --- | --- | --- | --- | --- | --- |
|  | Ch01 | Ch02 | Ch03 | Ch04 | Ch05 | Ch06 | Ch07 | Ch08 | Ch09 | Ch10 | Ch11 | Ch12 |
| Ch01 | 1 | 0.032 | 0.035 | 0.049 | 0.031 | 0 | 0.101 | 0.011 | 0 | 0.046 | 0.001 | 0 |
| Ch02 | 0.032 | 1 | 0.105 | 0.052 | 0.03 | 0 | 0.027 | 0.087 | 0 | 0.027 | 0.002 | 0.009 |
| Ch03 | 0 | 0.217 | 1 | 0.227 | 0.032 | 0 | 0.004 | 0.022 | 0 | 0.063 | 0.001 | 0.005 |
| Ch04 | 0 | 0.063 | 0.326 | 1 | 0.026 | 0 | 0.001 | 0.008 | 0 | 0.131 | 0.001 | 0.003 |
| Ch05 | 0 | 0.02 | 0.115 | 0.537 | 1 | 0 | 0.001 | 0.004 | 0 | 0.084 | 0.044 | 0.008 |
| Ch06 | 0.017 | 0.033 | 0.038 | 0.099 | 0.288 | 1 | 0.004 | 0.006 | 0 | 0.032 | 0.008 | 0.082 |
| Ch07 | 0.028 | 0.017 | 0.009 | 0.012 | 0.05 | 0 | 1 | 0.337 | 0.012 | 0.344 | 0.025 | 0.026 |
| Ch08 | 0 | 0.077 | 0.044 | 0.015 | 0.045 | 0 | 0.151 | 1 | 0.012 | 0.126 | 0.022 | 0.02 |
| Ch09 | 0 | 0.014 | 0.213 | 0.05 | 0.028 | 0 | 0.02 | 0.226 | 1 | 0.556 | 0.024 | 0.007 |
| Ch10 | 0 | 0.005 | 0.068 | 0.211 | 0.021 | 0 | 0.007 | 0.088 | 0.04 | 1 | 0.025 | 0.004 |
| Ch11 | 0 | 0.004 | 0.027 | 0.125 | 0.851 | 0 | 0.004 | 0.037 | 0.014 | 0.718 | 1 | 0.087 |
| Ch12 | 0 | 0.006 | 0.011 | 0.024 | 0.228 | 0 | 0.025 | 0.037 | 0.015 | 0.155 | 0.149 | 1 |

## Footnotes

1 https://github.com/theislab/deepflow, last accessed on 7th of May 2019

2 https://github.com/raghakot/keras-resnet, last accessed on 7th of May 2019

3 https://github.com/saeyslab/DeepLearning_for_ImagingFlowCytometry

4 http://github.com/saeyslab/cifconvert, last accessed on 7th of May 2019

5 https://github.com/saeyslab/DeepLearning_for_ImagingFlowCytometry

6 https://github.com/saeyslab/cifconvert

